# Quadrant darkfield (QDF) for label-free imaging of intracellular puncta

**DOI:** 10.1101/2024.08.05.606686

**Authors:** Tarek E. Moustafa, Rachel L. Belote, Edward R. Polanco, Robert L. Judson-Torres, Thomas A. Zangle

## Abstract

**Significance:** Measuring changes in cellular structure and organelles is crucial for understanding disease progression and cellular responses to treatments. A label-free imaging method can aid in advancing biomedical research and therapeutic strategies.

**Aim:** This study introduces a computational darkfield imaging approach named quadrant darkfield (QDF) to separate smaller cellular features from large structures, enabling label-free imaging of cell organelles and structures in living cells.

**Approach:** Using a programmable LED array as illumination source, we vary the direction of illumination to encode additional information about the feature size within cells. This is possible due to the varying level of directional scattering produced by features based on their sizes relative to the wavelength of light used.

**Results:** QDF successfully resolved small cellular features without interference from larger structures. QDF signal is more consistent during cell shape changes than traditional darkfield. QDF signals correlate with flow cytometry side scatter measurements, effectively differentiating cells by organelle content.

**Conclusions:** QDF imaging enhances the study of subcellular structures in living cells, offering improved quantification of organelle content compared to darkfield without labels. This method can be simultaneously performed with other techniques such as quantitative phase imaging to generate a multidimensional picture of living cells in real-time.

## 1. Introduction

Measuring dynamic reorganization in cellular structure and organelles is vital for the study of disease progression and response to treatment^1-3^. As cells switch phenotypes in response to environmental or genetic signals, corresponding changes are observed in cellular components including mitochondria^4^, endoplasmic reticulum^5, 6^, Golgi apparatus^7^, melanosomes^8, 9^, microtubules^10-12^ and the plasma membrane^13-15^. These changes can be related to cellular events such as necrosis^16^, senescence^17^, and the emergence of drug resistance^8^. The quantification of intracellular dynamics, therefore, serves as a critical indicator of a cell’s health, behavior, and its response to therapeutic interventions.

Imaging approaches are commonly used to measure changes in subcellular structure. For example, electron microscopy (EM) is commonly used to study details of organelle structure in fixed cells^8^ but cannot quantify changes in living cells in real time. Fluorescence microscopy has been widely used to study cell structure, including measurement of damage to plasma membranes^15^, measurements of melanosome maturation^18^, localization of nuclei^19^, and tracking lysosomes during autophagy and other cell states^20, 21^. However, the need for a fluorescent molecule that binds to the specific target or for a cell to express a fluorescently tagged protein is one of the limiting factors of fluorescence microscopy. Additionally, labels can affect cell behavior and the need for high intensity illumination can cause phototoxicity especially during live cell imaging^22^.

Intrinsic scattering and autofluorescence can also be used for label-free imaging of cell structure. Side scatter measurements in flow cytometry can be used to quantify changes in subcellular structure without the need for fluorescent labels^23^. Whereas forward scatter measurements predominantly measure cell size, side scatter reflects the granularity of internal contents of the cell^23^. However, flow cytometry cannot track changes in the same cell over time^24^. Raman spectroscopy utilizes the weak inelastic scattering of light that is dependent on molecular composition to identify organelles and molecular composition within the cell, such as nucleic acids, mitochondria, and endoplasmic reticulum^25^. However, Raman signal is typically weak, necessitating the use of high powered lasers or long exposure times to produce sufficient signal to noise ratios^25^. Light scattering spectroscopy (LSS) uses elastic backscattering that is dependent on both wavelength and particle size to measure the concentration and size of organelles^26-30^. For example, the ability of LSS to measure the size of organelles and changes in structure allowed for the detection of precancerous and malignant cells in multiple cancer types^26, 28, 29^. However, LSS is not an imaging technique and so cannot be used to study the localization of organelles inside cells. Autofluorescence is the natural phenomena of proteins emitting light when excited using a suitable wavelength without the use of labels^31^. However, the main limitations of autofluorescence are the low signal that organelles produce, need for a specific wavelength of light for each organelle, and the limited number of organelles and structures that naturally fluoresce^31^.

Computational microscopy provides methods to measure the intrinsic contrast caused by refractive index variation within cells. Multiple organelles, including lysosomes and mitochondria, differ slightly in density and refractive index from the surrounding cytoplasm^32, 33^. This difference can be quantified using quantitative phase imaging (QPI)^34^ which measures phase shift as light passes through the cell^35, 36^. One recent example used QPI to track lysosomes in living cells without use of fluorescent tags^32^. However, tracking organelles can be difficult with two-dimensional QPI due to the signal from overlapping cell components. Three-dimensional computational methods such as Fourier ptychography^37^ or three-dimensional differential phase contrast^38^ provide the needed information to resolve density or refractive index changes in each 3D voxel within the cells. However, these techniques require tens of images per field of view, limiting their temporal resolution and their ability to conduct high throughput experiment in multi well plates.

Some of the simplest techniques to study changes in cellular structure are brightfield and darkfield imaging. Brightfield signal correlates with absorption which is usable in studying organelles such as melanosomes that absorbs light^39^, but limited for other cell structures. On the other hand, many subcellular features scatter light due to variation in refractive index between organelles and cytoplasm^40^. For cellular organelles (0.1-10 μm) imaged using visible light (380-700 nm), this scattering is explained using Mie scattering theory^23, 41^ and features within this size range can be detected using darkfield imaging. Early uses of darkfield were to detect contaminants in blood samples and to differentiate cell types due to its inherent contrast even between objects close to the refractive index of the media used^42^. Since then, darkfield has been used to quantitatively measure the size of red blood cells^43^, to track in real time respiratory syncytial virus (RSV) infecting cells using gold nanoparticles^44^, to measure nanoparticle distribution in lung cells^45^, and to visualize and count sub-micron particles in suspension^46^. When performing darkfield using transillumination, larger features, such as the boundaries of cells tend to have higher signal. This is due to more directional refraction of light into the objective from large features^41, 47, 48^. As the feature size gets smaller, light starts to scatter in a cone, bending more light away from the objective and decreasing signal. For a given wavelength, this cone keeps growing as the feature size gets smaller leaving only a negligible amount of light to be collected by the objective. This change is commonly used for sizing particles^47-49^. Additionally, the difference in directionality between larger and smaller objects means that illumination from different directions can potentially resolve different features based on size during imaging.

In this work, we develop a computational darkfield approach we call quadrant darkfield (QDF). QDF differentially resolves smaller cellular features which scatter broadly as explained by Mie scattering (∼ λ) from larger features that more uniformly refract light (>> λ). This allows imaging of small features of the cell such as cellular structures and organelles without interference from larger structures such as cell edges. We demonstrate QDF on an inverted microscope using a programmable LED array as the light source. This approach can be used simultaneously with other modalities such as QPI to collect multimodal images of cells in standard multi well plates. Here, we demonstrate QDF’s ability to resolve features in polystyrene beads and both pigmented melanoma cells and non-pigmented breast cancer cells. We demonstrate that QDF is not impacted by changes in shape such as occur during cell division, in contrast to traditional darkfield which is highly dependent on cell shape. Finally, we show concordance between QDF signal and side scatter from flow cytometry to differentiate melanoma cells with different levels of pigmentation.

## Materials and Methods

### 2.1. Darkfield and QDF data acquisition

QDF was performed on a custom-built microscope consisting of a 0.25 NA, 10x objective (PLN 10X, Olympus, Japan), a monochrome 1920 × 1200 CMOS camera (GS3-U3-23S6M-C, Teledyne FLIR, USA) and a 180 mm tube lens (Fig. 1a)^50^. The LED array was illuminated between 1.05x to 1.33x NA_objective_ (Fig. 1b). Four images were captured using quadrant darkfield illumination patterns (Fig. 1c) at an exposure time of 220 ms and a gain of 25 dB. Darkfield images were computed by summing the four quadrant images. For QDF processing, the quadrant images were combined to compute the edge image, *E*, as:

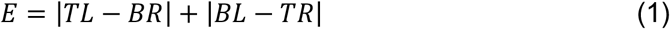

where *TL, BR, BL*, and *TR* refer to images captured under top-left, bottom-right, bottom-left, and top-right illumination, respectively (fig. 1b). All operations are performed pixel-wise. The edge image was then subtracted from a scaled darkfield image to produce the QDF image:

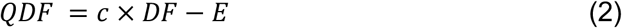

where *c* is a scaling factor that is system specific and determined experimentally to match the darkfield signal at the edge of objects with the edge image and is typically in the range 0.8-1.0 depending on the experimental setup.

**Figure 1.**
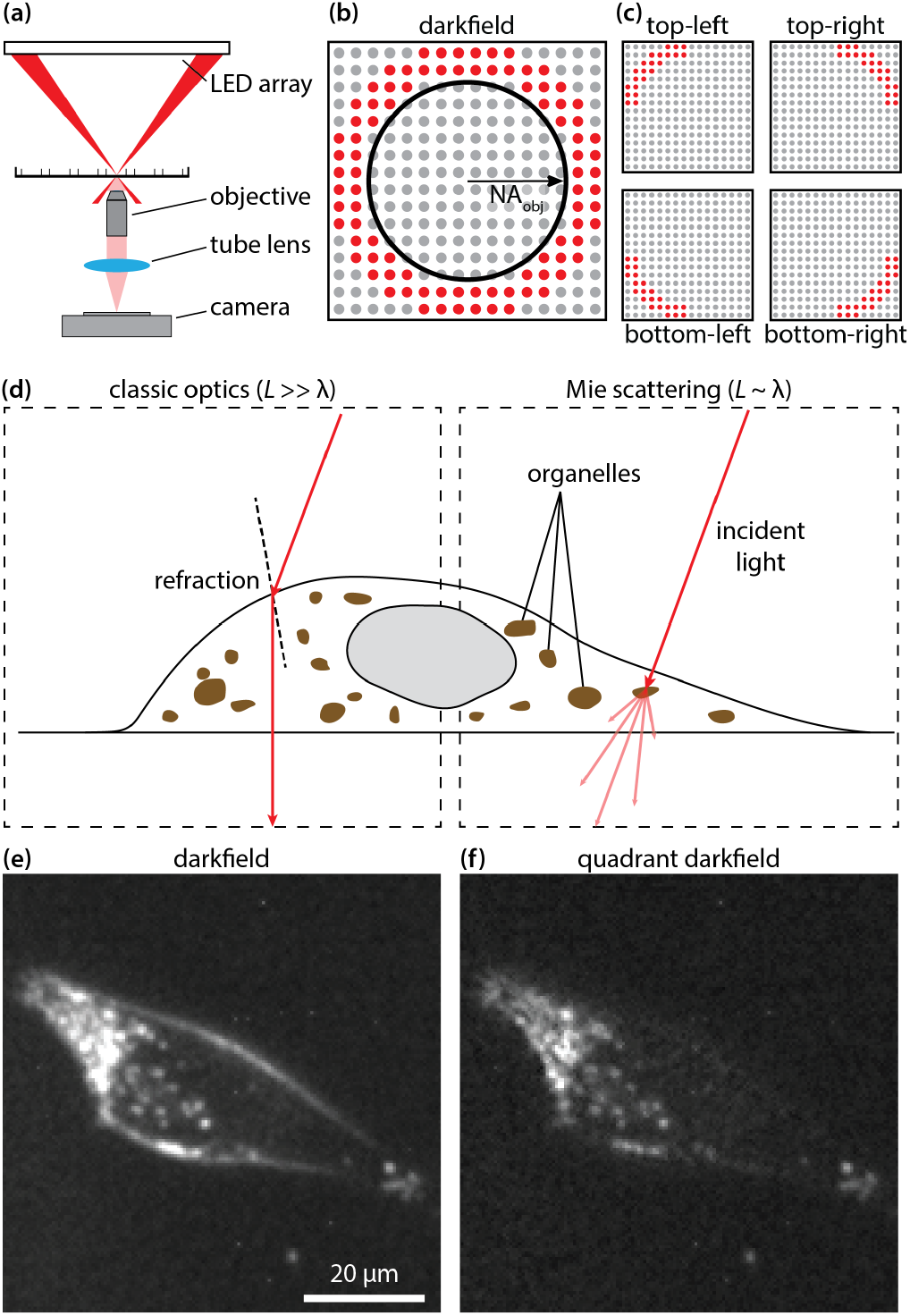
(a) Schematic diagram of an inverted microscope with an LED array as a light source in darkfield illumination setup. (b) Bottom-up view of the LED array illumination pattern in darkfield with the numerical aperture of the objective superimposed. (c) The illumination pattern used for each of the quadrants in QDF. (d) Diagram show how light is directionally refracted by large features according to Snell’s law (left) relative to how light is scattered when hitting a subcellular feature according to Mie scattering theory (right). (e) Darkfield image of a melanoma PDX cell (MTG021). (f) QDF image of the same cell from e.

### 2.2. QPI data acquisition

QPI was obtained using differential phase contrast (DPC) and reconstructed using Tikhonov deconvolution^51, 52^. DPC images were captured with an exposure time of 50 ms, a gain of 25 dB, coherence parameter of 1.25, and a regularization parameter of 4×10^−3^, which was determined experimentally as described previously^50^.

### 2.3. Cell culture

#### 2.3.1. MDA-MB-231

MDA-MB-231 were acquired from ATCC (HTB-26, ATCC, USA) and passaged on 100 mm plates (12-567-650, Thermo Fisher Scientific, USA) in RPMI (11875093, Thermo Fisher Scientific) with 10% FBS (10-437-028, Thermo Fisher Scientific) and 1% Penicillin-Streptomycin (15140122, Thermo Fisher Scientific). For passaging, cells were washed with Dulbecco’s phosphate buffered saline (14190144, Thermo Fisher Scientific) then incubated with Trypsin (15400054, Thermo Fisher Scientific) at 37 °C and 5% CO_2_ for 7 min followed by trypsin deactivation using RPMI medium at 1:1 ratio. The dissociated cell suspension was centrifuged at 400 x*g* at room temperature for 4 minutes before resuspension in RPMI medium and splitting at a 1:4 ratio.

#### 2.3.2. MTG084 and MTG021

MTG084 (AM084) and MTG021 (ASM021) cells were previously generated from patient derived xenograft (PDX) models of melanoma (Huntsman Cancer Institute, Preclinical Research Resource)^53^ and grown in AM3 media (80% MCDB153 (#M7403, Sigma, USA), 20% L-15 media (#11415-064, Thermo Fisher Scientific), 2.5% FBS (#FB5001-H, Thermo Fisher Scientific), 1X Insulin-Transferrin-Selenium X (#51500-056, Thermo Fisher Scientific,), 5 ng/mL EGF, 0.2% BPE, 10 ng/mL Insulin like growth factor, 5 µg/mL transferrin, 3 ng/mL BFGF, 3 µg/mL heparin, 0.18 µg/mL hydrocortisone, 10 nM endothelin 1, 1.68 mM CaCl2) in 5% CO_2_ at 37°C.

### 2.4. Image acquisition and processing

#### 2.4.1. Image acquisition

Cells were plated at 12,500 cells in each well of a 24 well plate (7000674, Greiner Bio-One, Germany) for imaging. Cells were incubated for 24 h (MDA-MB-231) or 48h (MTG021 and MTG084) prior to imaging and placed inside the microscope incubator for 30-45 min prior to imaging. Cells were imaged every 20 min with a single autofocus between imaging cycles to account for thermal and z-stage drift. Nine imaging positions were chosen at the center of each well to avoid scattering from the edges of the well.

#### 2.4.2. QPI processing

QPI images were background corrected by masking cells and fitting an 8th order polynomial to background pixels that was subtracted from the raw phase image. Single cells were segmented using a watershed algorithm and cell dry mass was computed using a cell average specific refractive increment of 1.8 × 10^−4^ m^3^/kg^35^. Segmented cells were tracked over time using the Crocker-Grier algorithm^54^ to aid in the detection of debris.

#### 2.4.3. Darkfield image processing

Darkfield images were scaled down to 12 bits by dividing by 16 and rounding to nearest integer to match the camera bit depth. An image from an empty reference position was subtracted from darkfield images to remove background signal. An 8th order polynomial fit was removed from the background of masked quadrant, and edge images.

### 2.5. Flow cytometry

Cells were harvested using 0.05% trypsin (#25300054, Thermo Fisher Scientific) followed by trypsin deactivation with 1:1 Soybean Trypsin Inhibitor (#17075029, Thermo Fisher Scientific). The dissociated cell suspension was centrifuged at 500 g, 4 °C, for 4 min, and resuspended in ice cold FACS buffer (0.1% BSA, 2.5% HEPES, in HBSS). Single-cell suspensions were counted, diluted to 1 × 10^6^ cells per 300 – 500 µl in ice-cold FACS buffer and passed through a 35 µm filter. Flow cytometry was performed using a BD FACS Aria sorter (BD, USA), BD Fortessa analyzer (BD, USA), and SONY SH800 sorter (SONY, Japan). SSC (Aria and Fortessa)/ BSC (SONY) analysis was conducted on single cells by gating to exclude debris and doublets using a two-step doublet discrimination gating strategy first with FSC-A vs FSC-H followed by SSC-A vs SSC-H or BSC-A vs BSC-H.

### 2.6. Statistics

Segmented objects were filtered using thresholds of area, track length, and averages of phase shift, darkfield and QDF to remove debris. Segmented cells were manually inspected to remove under segmentation of multiple cells and over segmentation of single cells. A least-squares linear regression was applied to QDF and darkfield signals against mass per area. Linear fits were compared to a no fit option of a flat line parallel to the *y*-axis using an *F*-test with a *p* value of 0.05. Correlation between variables was computed using the Pearson correlation coefficient in Matlab (Matlabs R2021a, Mathworks, USA).

### 2.7. Signal to noise ratio calculation

To quantify signal, a threshold mask equal to 4x the 99^th^ percentile of the background signal in each image was applied to each QDF image. The intersection of single cell labels and the threshold mask is the QDF signal from puncta in each cell. The noise level is defined as the standard deviation of background signal outside masked cells. Results are reported as mean +/-standard deviation.

### 2.8. Bead sample preparation

20 µm Polystyrene beads (18329-5, Polysciences, USA) were embedded in NOA73 (NOA73, Norland Products, USA) sandwiched between a standard microscope slide (12-544-4, Fisher scientific, USA) and a no. 1.5 cover glass (22-037-082, Fisher scientific) and cured under ultraviolet radiation (IntelliRay, Uvitron, USA).

## 3. Results

Using an LED array as illumination source (fig.1a) enables use of different illumination schemes, including darkfield imaging with a ring of LEDs outside the numerical aperture of the objective lens (fig. 1b)^51^. In QDF, this darkfield illumination ring is divided into four quadrants (fig. 1c), with one image acquired under illumination from each quadrant to capture illumination-angle specific darkfield images. For living cells, there are two main types of scattering based on the relative size of the cellular feature to the wavelength of light. For features much larger than the wavelength *e*.*g*. the outer cell membrane, the path of light can be predicted by classical optics (Snell’s Law, fig. 1d). For features close in size to the wavelength *e*.*g*. sub-cellular puncta or organelles, the path of light can be explained by Mie scattering theory (fig. 1d). Features of both sizes appear in darkfield (fig. 1e). However, the larger darkfield features (e.g. cell boundaries) often obscure smaller features (*e*.*g*. organelles), especially near cell edges. QDF separates the darkfield signal based on directionality of light allowing for a cleaner image of sub-cellular features (fig. 1f).

To validate feature differentiation with QDF, we embedded 20 µm polystyrene beads into optical adhesive for use as an imaging phantom. When imaged in darkfield, these beads behave as spherical lenes with the center rays passing straight through and the edges refracting light, resulting in beads appearing as a bright ring (fig. 2a). In most beads, some weaker signal inside and outside the bead due to diffraction around the bead as well as light from other out-of-focus sections of the bead is also observed (fig. 2b). Under asymmetric quadrant illumination, we observe light being refracted by the opposing quarter of the bead (fig. 2c-d). When we combine the quadrants into an edge image (eq. 1), the edges of the bead are captured clearly (fig. 2e). Subtraction of this edge image in QDF removes nearly all the signal from most beads (fig. 2f). We also observed a number of beads with imperfections, possibly due to material deformations or the accumulation of impurities as the beads dry on top of the glass slide^55^. Looking closer at an imperfect bead (fig. 2g) darkfield imaging shows both the imperfection as well as the bead edge. Under asymmetric quadrant illumination, the imperfection is always present in all images along with the opposing edge (fig. 2h,i). This validates that the signal from the smaller features is less directionally dependent on the angle of illumination in darkfield when compared with signal from the edges. Thus, when the quadrant images are combined (eq. 1), the resulting edge image primarily has signal from the edges. When the QDF image is computed, the imperfections at the center of the bead are observed clearly without being obscured by signal from the edges (fig. 2k).

**Figure 2.**
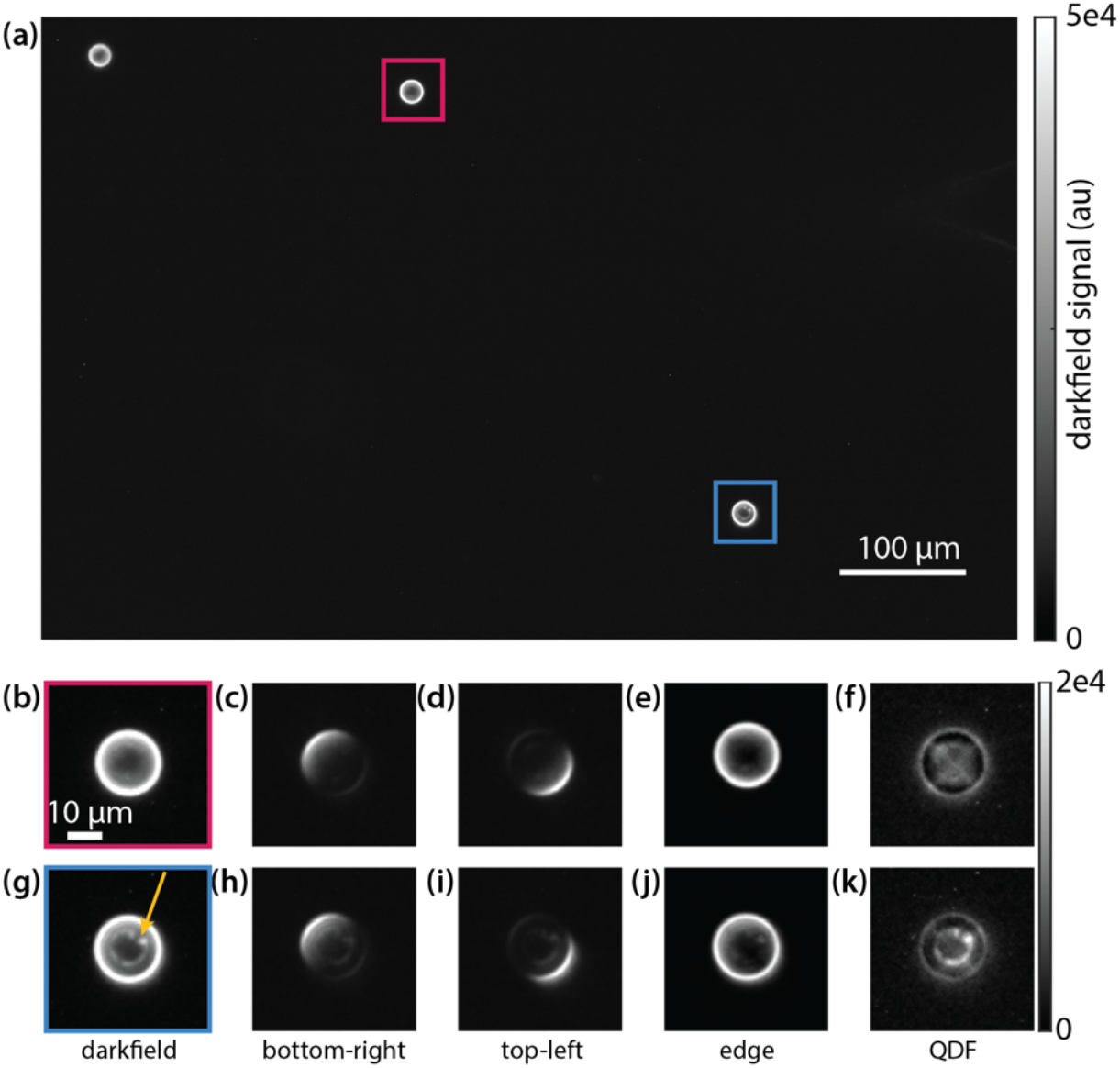
(a) Full field of view image of 20 µm polystyrene beads in darkfield. (b) Zoomed in darkfield image of a clean bead. (c) Top-left illuminated quadrant image of the clean bead. (d) Bottom-right illuminated quadrant image of the clean bead. (e) The resulting edge image from combining the four quadrants. (f) QDF image of the clean bead. (g) Zoomed-in image of the imperfect bead. Arrow highlights visible imperfection. (h) Top-left illuminated quadrant image of the imperfect bead. (i) Bottom-right illuminated quadrant image of the imperfect bead. (j) The resulting edge image from combining the four quadrants. (k) QDF image of the imperfect bead. Red and blue boxes in (a) correspond to fields of view in (b-f) and (g-k), respectively.

Next, we applied QDF to a melanoma cell line, as they have abundance of melanosomes, pigment producing lysosome related organelles, with a size range between ∼100 nm to 1 µm^9, 56^. Within this size range, Mie theory predicts that QDF will be able to separate subcellular features from cell edges due to the broad directionality of scattering from organelles relative to the directional uniform scattering at cell boundaries. We additionally imaged the cells using quantitative phase imaging (QPI) to aid in segmenting the cells (fig. 3a). In darkfield, cells show varied signal based on shape and cellular content with more rounded cells producing higher signal from the edges (fig. 3b). Using QDF, we were able to differentiate smaller features within cells from cell edges giving a clear view of puncta inside the cells. QPI also quantifies the dry mass of cells and helps differentiate flat from more rounded cells based on the density at each pixel^57^ (fig. 3d). When we view two cells with high mass density (reflecting cell rounding) and low mass density (reflecting a flatter cell morphology) in darkfield, the difference in signal from cell edges is apparent (fig. 3e). In these same cells, the edge image (eq. 1) shows the edge of each cell. (fig. 3f). The resulting QDF image shows much greater contrast in intracellular puncta (fig. 3g). Additionally, we characterized the signal to noise ratio (SNR) of individual puncta resolved by QDF in MTG021 as 31 ± 6 (fig. S1). In MTG084, an additional melanoma cell line with different pigmentation, the QDF SNR was 24 ± 5.

**Figure 3.**
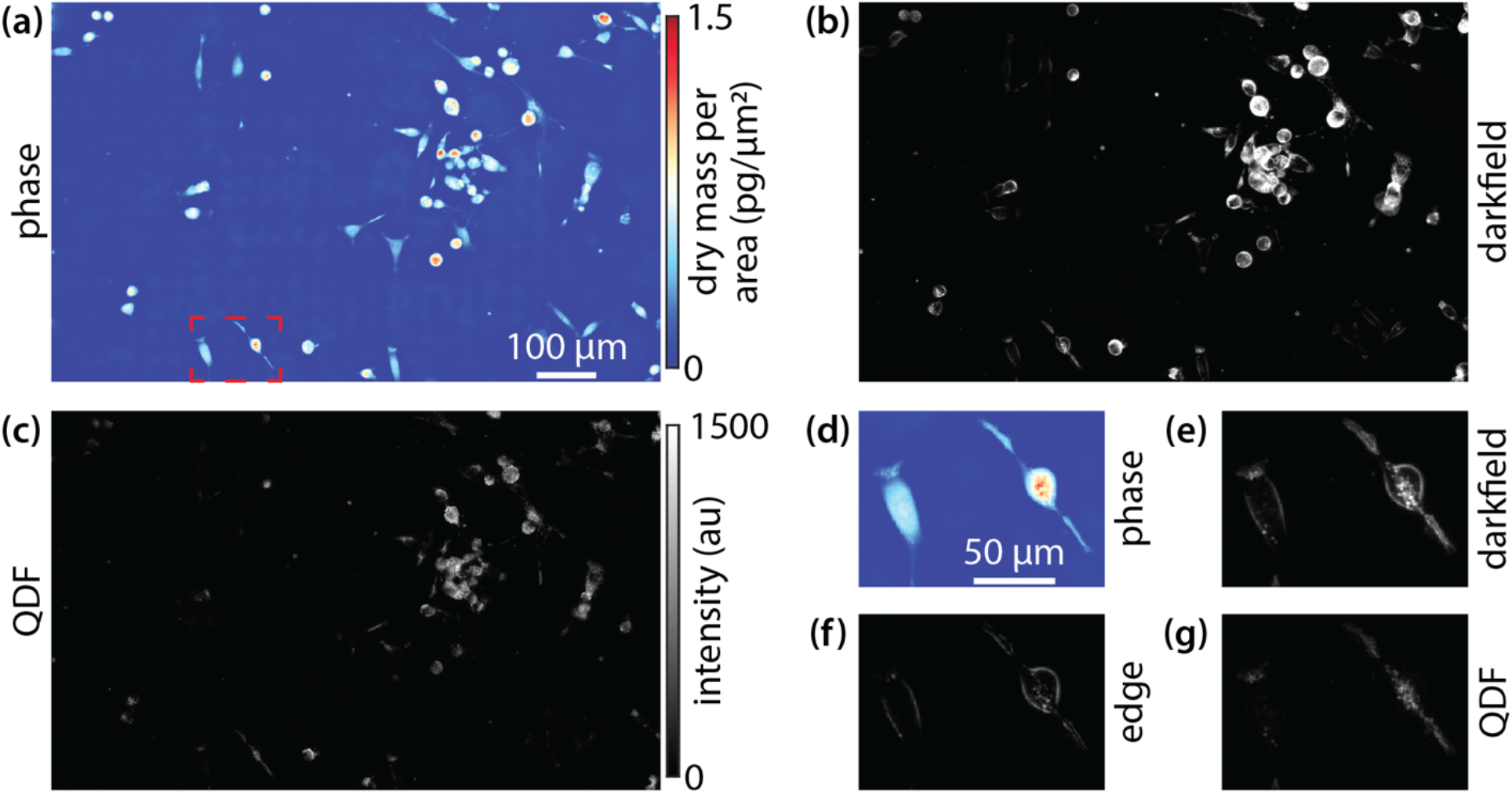
Melanoma cell line MTG021 imaged via (a) QPI, (b) darkfield, and (c) QDF. (d) QPI of two adjacent cells of varying shape and mass density from the outlined region in (a). (e) Darkfield image of these cells. (f) Edge image resulting from the quadrant image processing. (g) Resulting QDF image of the two cells.

One of the issues with using traditional darkfield imaging for longitudinal tracking of subcellular features is the dependency of the signal from cell edges on the shape of the cell. Cells change morphology naturally throughout the cell cycle. For example, adherent cells become rounded just prior to cell division^58, 59^. To validate QDF signal independence from shape, we imaged a breast cancer cell line that does not contain melanosomes (fig. 4a).

**Figure 4.**
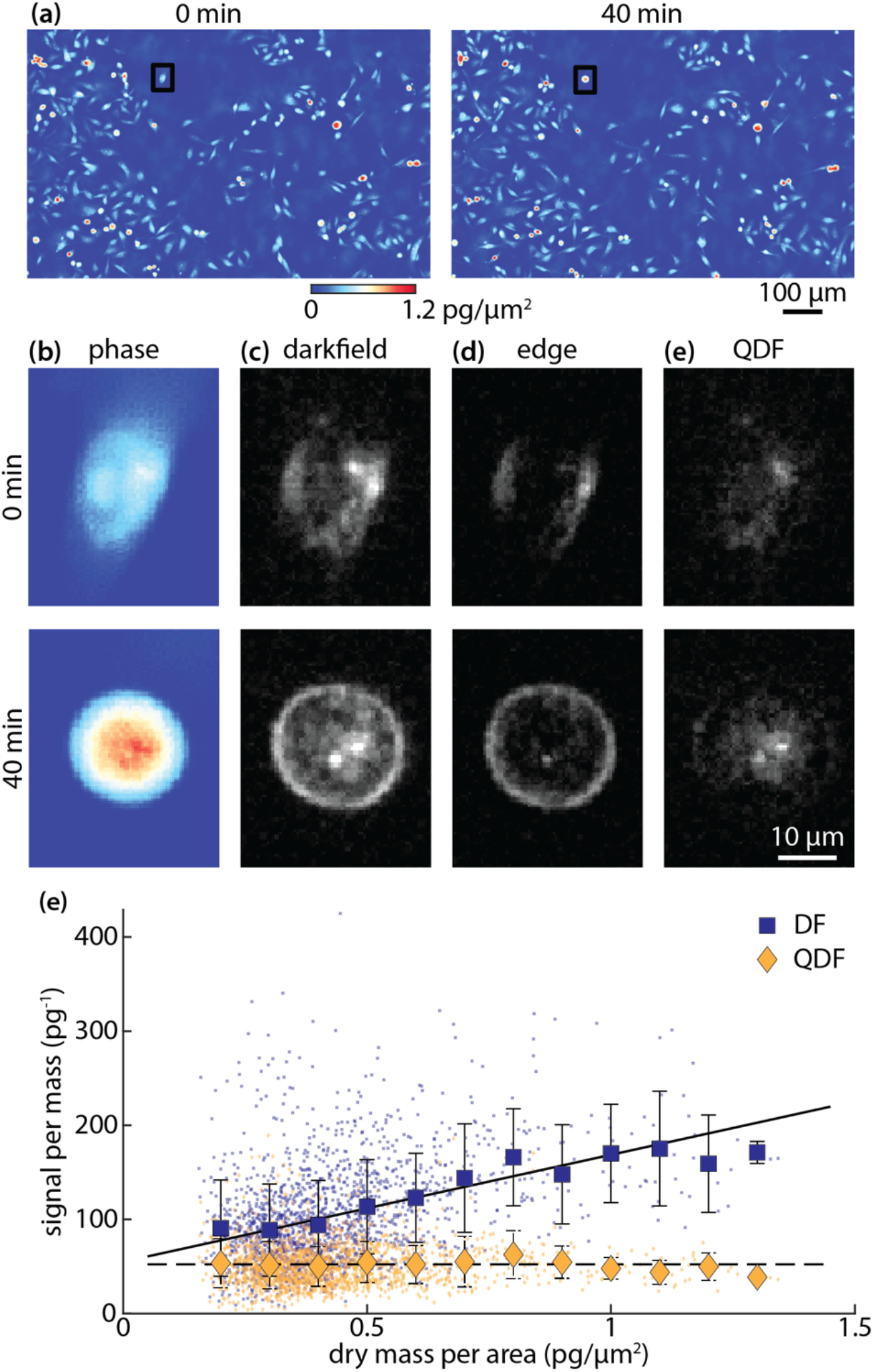
(a) phase images of MDA-MB-231 breast cancer cells (b) darkfield images (c) edge images (d) QDF images. Sub-images in (b-e) correspond to black boxed cell in (a). (e) scatter plot of darkfield (dark blue) and QDF (dark yellow) per mass (y-axis) against mass per area (x-axis) as a proxy of shape change. Larger points are the average within bins of 0.1 pg/µm^2^. Error bars show standard deviation within each bin. Lines show linear fits to the data as an indication of the overall trend (solid: darkfield, DF; dashed: QDF).

As an example of the shape changes that occur during cell cycle progression, one individual cell changes shape from flat to rounded over 40 min (fig 4b). In darkfield, the signal increases significantly due to the additional lensing effect of the rounded edges of the cell (fig. 4c). By looking at the computed edge images (eq. 1), the difference in the brightness of the edges is drastic (fig. 4d). In the resulting QDF images, the density of the signal increases more moderately as the puncta are pulled closer to each other (fig 4e). During this transition, the total darkfield signal increased by 40%. On the other hand, the total QDF signal change was 5%.

This is within the accuracy limits of our system, reflecting that the total organelle content of the cell did not appreciably change over this period. When analyzing the QDF and darkfield signal on a population basis, the dependency of darkfield on shape, as measured using mass per area^57^, is clear (fig S2). QDF does show a weak dependency on mass per area, potentially due to an increase in mass resulting in an increase in scattering organelle content. This is confirmed by looking at darkfield and QDF signal per mass against mass per area (fig 4e). These data indicate that darkfield signal is significantly more dependent on cell shape than QDF (Pearson’s correlation coefficient, *R*, for darkfield = 0.39 and for QDF = 0). When plotting QDF against mass (fig S3), QDF shows a strong dependency on mass (*R*_*QDF*_ = 0.60). However, this dependency is not perfectly linear.

Finally, we examined the use of QDF to differentiate pigmentation in melanoma cell lines with varying levels of pigmentation (fig. 5a). Flow cytometry was able to differentiate the two cell lines based on side scatter (fig. 5b, fig. S4). In QDF images of the two cell lines, the difference in puncta can clearly be seen (fig 5c). Total QDF was also able to differentiate the two different cell lines (fig 5d), similar to flow cytometry. On the other hand, darkfield shows significantly more overlap between the two cell lines when compared to QDF (fig. S5) and cannot localize puncta as clearly as QDF (fig.1e,f, fig. S6).

**Figure 5.**
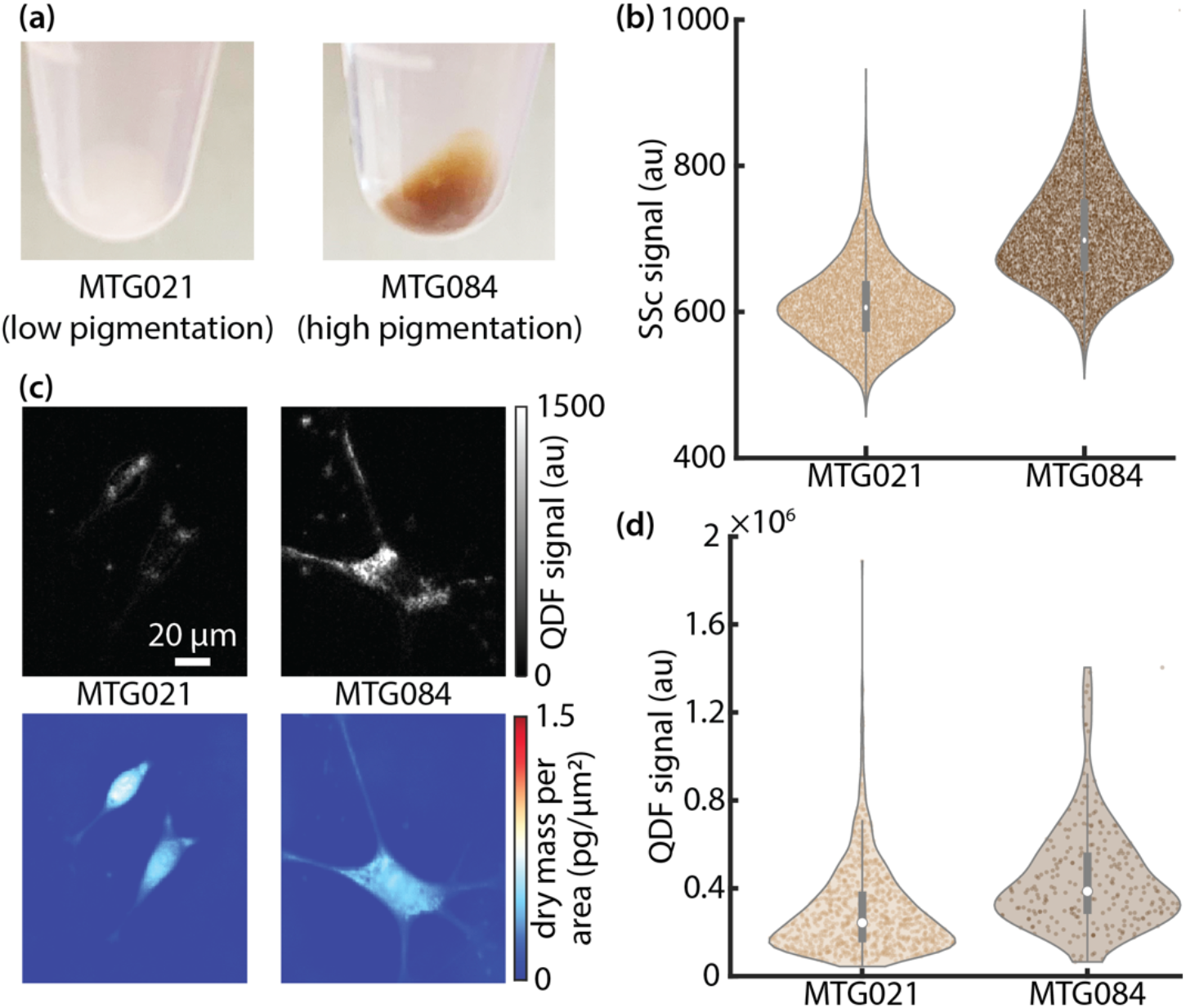
(a) Cell pellets of two melanoma cell lines with significant difference in pigmentation. (b) Distributions of side scatter measurements from flow cytometry for the two cell lines. (c) QDF and QPI images of example cells for each cell line. (d) Distributions of QDF measurements of the two cell lines.

## 4. Discussion

Here, we demonstrated QDF as a label-free technique for imaging sub-cellular puncta. Unlike darkfield, QDF signal does not depend on cell shape providing a more reliable quantitative measurement of sub cellular structure. Additionally, the edge measurement used to compute QDF images can be used as a fast indicator for shape change to signal stress or cell cycle event like division. This can potentially aid in better cell lineage tracking over time. We also demonstrated that QDF can be used to measure pigment-containing melanosomes within melanoma cell lines, with correlation to flow cytometry.

However, it should be noted that QDF shares some limitations with darkfield. These limitations include the need for alignment to avoid stray light, use of a bright illumination source for increased SNR, and imaging of thin samples to avoid scattering from out-of-focus layers. QDF is additionally diffraction limited, just as for conventional darkfield or brightfield imaging. Additionally, based on Mie theory the minimum size of observable particles with QDF is approximately one fifth of the wavelength of light used due to sharp drop in scattering signal beyond this point^41, 60^.

As a computational imaging technique, QDF is relatively low to moderate in complexity. In contrast to 3D ptychographic methods which require hundreds of images to be captured^38, 61, 62^, QDF differentiates sub-cellular structures using four images. This enables the use of QDF for applications requiring high throughput, such as drug screening applications^63^.

We showed that QDF was able to distinguish puncta inside cells of different masses (fig. 3b,e) and from different origins including a breast cancer cell line and non-pigmented melanoma line (figs. 4d, 5c). The ability of QDF to monitor subcellular heterogeneity within a single population (figs. 4e, 5d) is of great potential as an indicator of emerging resistance in melanoma^8^. QDF signal in more mature cells showed brighter puncta which can potentially be used to track melanosome maturation^9^. Additionally, QDF can be run simultaneously with QPI to increase the dimensionality of the data which can improve phenotype-based drug response assays^63^.

Compared to flow cytometry, QDF is inherently an imaging technique that produces detailed images of cells. These images can be used to study sub cellular dynamics over time^64^. QDF can be integrated with other modalities, including fluorescence imaging, for better characterization of cellular puncta.

## Acknowledgements

This work was supported by grants from NIH NCI (1R01CA276653, TAZ and RLJT), DoD CDMRP (W81XWH2210495, RLB), and the University of Utah (1U4U, TAZ and RLJT). We utilized the Huntsman Cancer Institute Shared Resource for Preclinical Research Resource and funding from the Cell Response and Regulation Program (TAZ and RLJT), both supported by the National Cancer Institute of the National Institutes of Health under Award Number P30CA042014.

## Code and Data Availability

Data and code are publicly available to recreate the findings of this article. Associated data is available at doi:10.5281/zenodo.13227201. Code is available at github.com/Zangle-Lab/QDF.

