## Supplementary Information for "Quadrant darkfield (QDF) for label-free imaging of intracellular puncta"

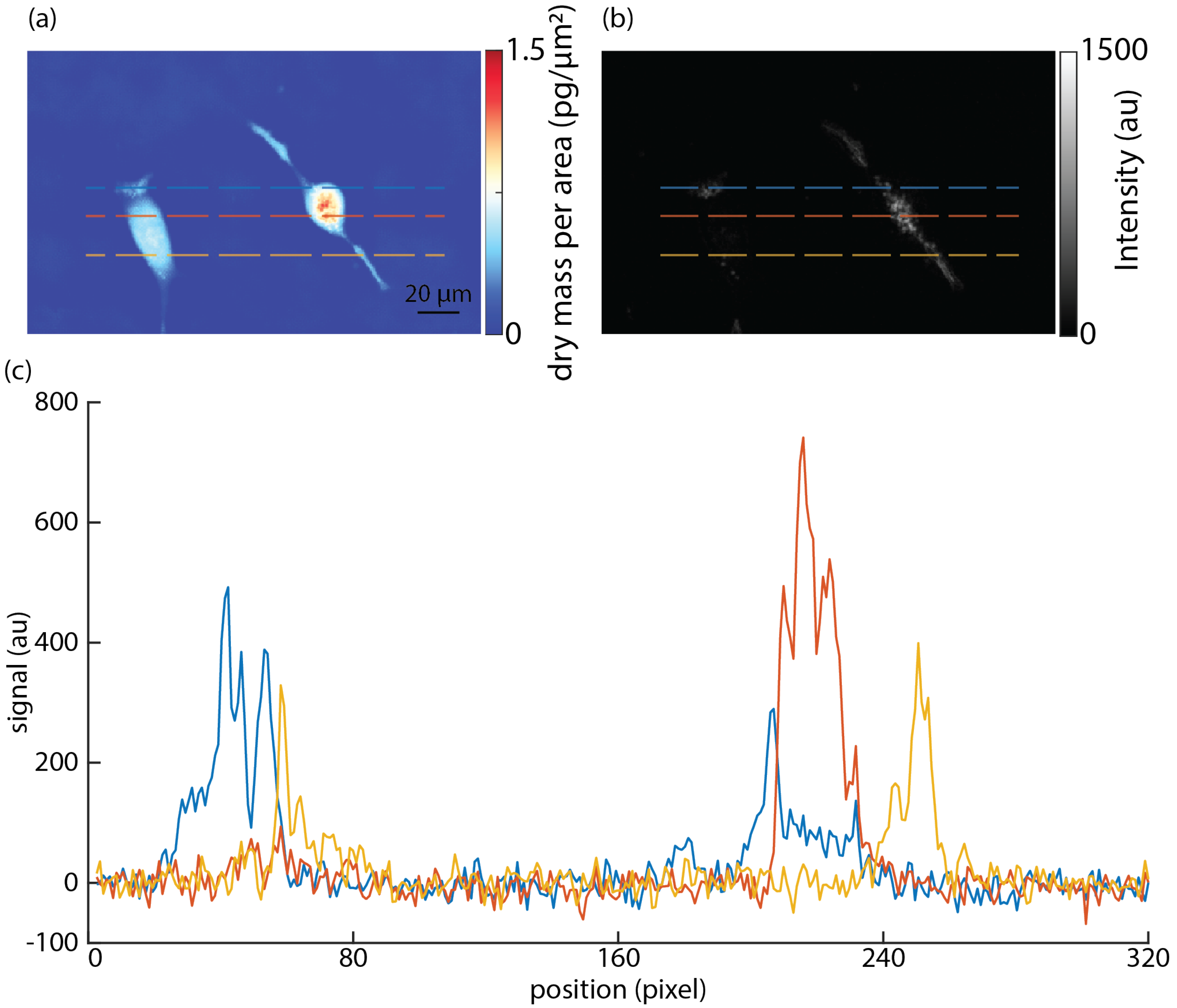


Fig S1 Zoomed view of MTG021 cells using phase (a) and QDF (b). (c) plotted line of QDF signal showcasing difference in signal between puncta and background as well as different puncta in each cell. Color of overlaid lines in (a) and (b) correspond to the color of plotted lines in (c) respectively.

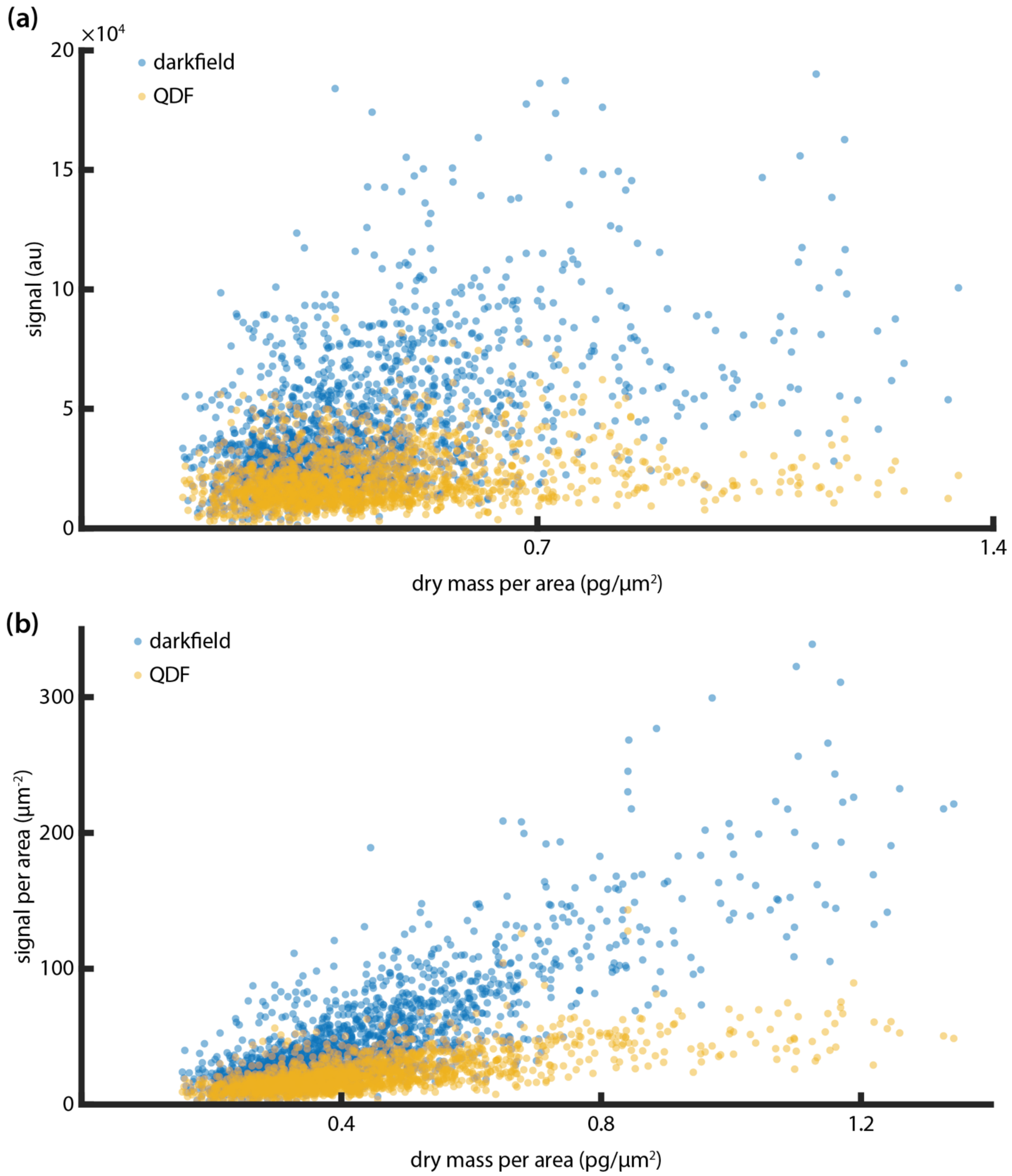


Fig S2 (a) scatter plot of darkfield (blue) and QDF (yellow) signal against dry mass per area as a proxy of a shape change (b) scatter plot of darkfield (blue) and QDF (yellow) per area signal against dry mass per area as a proxy of a shape change.

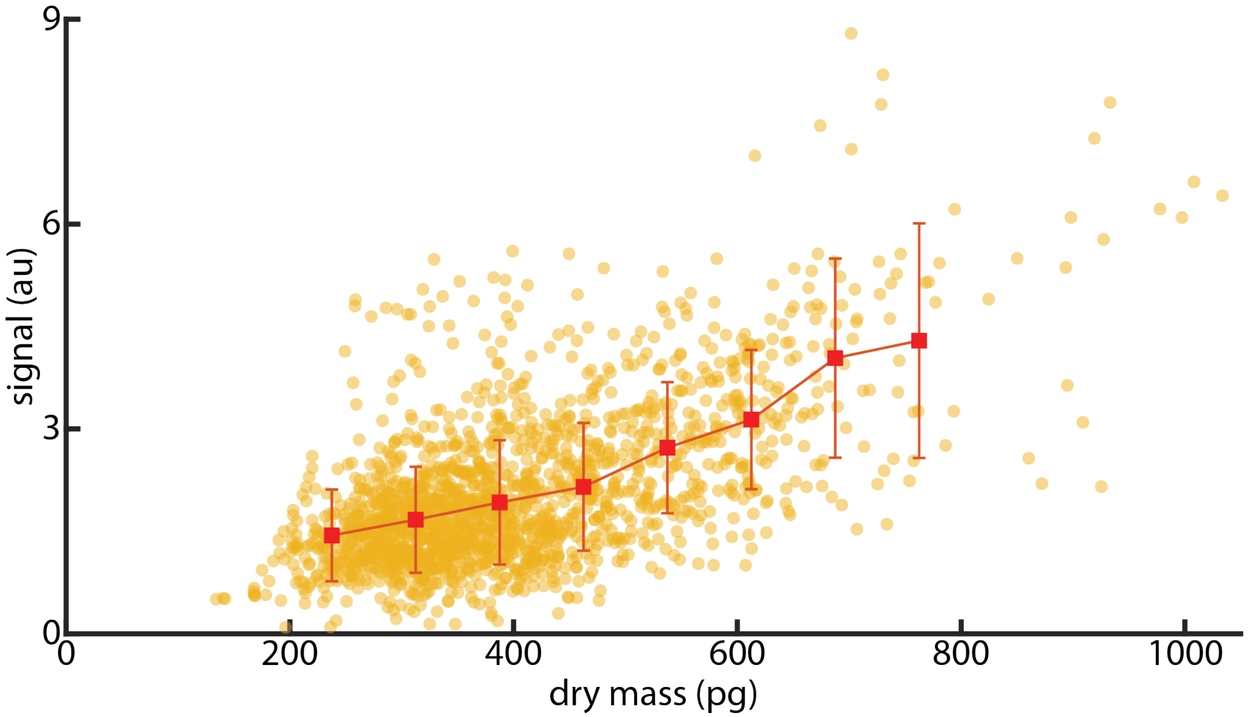


Fig S3 scatter plot of QDF (yellow) signal against dry mass. Data points are binned together (red marker) every 75 pg from 200 pg to 800 pg. Y-values of red markers are the mean of the data points within the bin edges. Error bars are the standard deviation of the binned data.

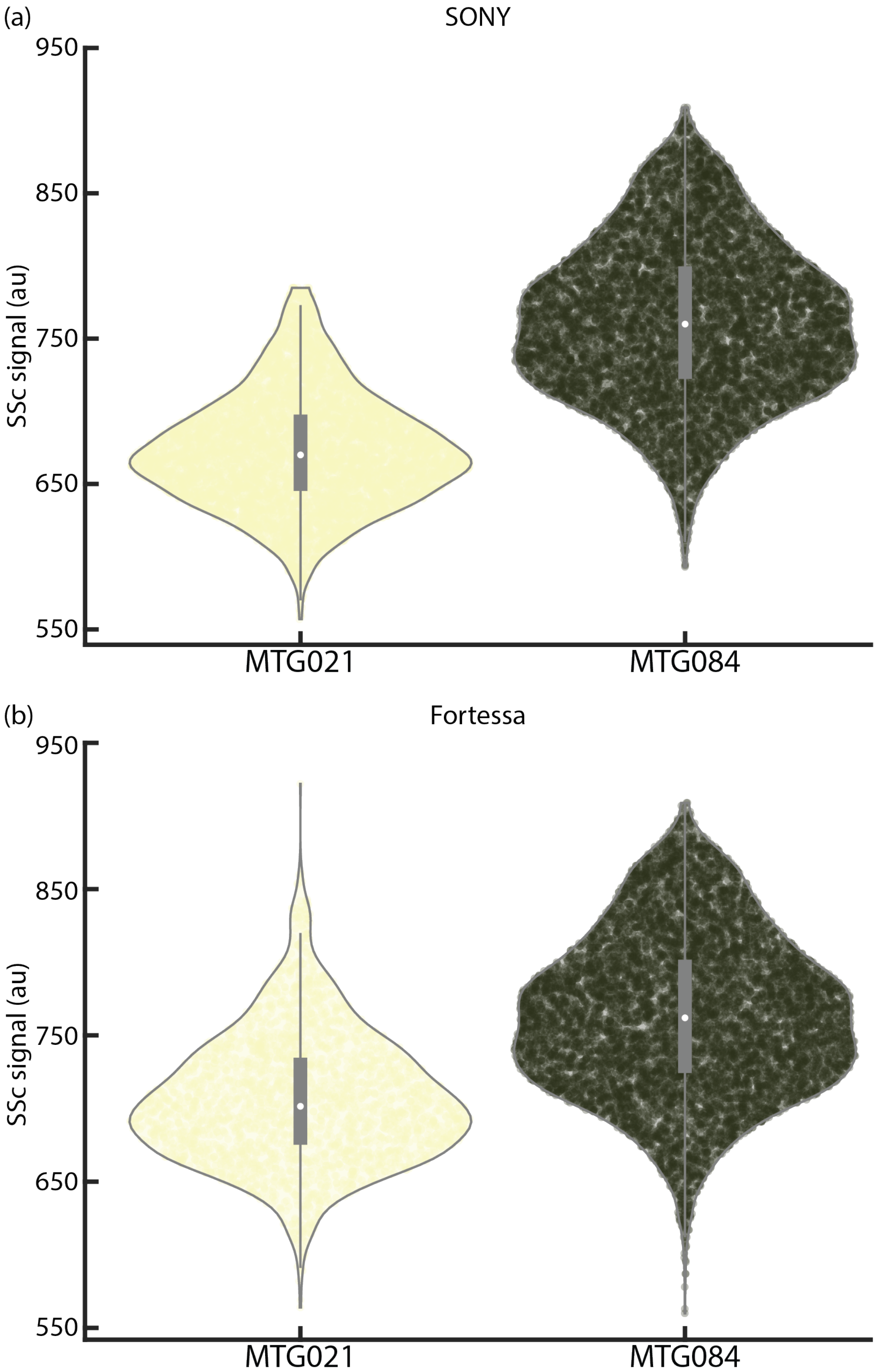


Fig S4. Distribution of side scatter signal for MTG021 and MTG084 from two flow cytometry devices (a) SONY SH800 and (b) BD Fortessa analyzer.

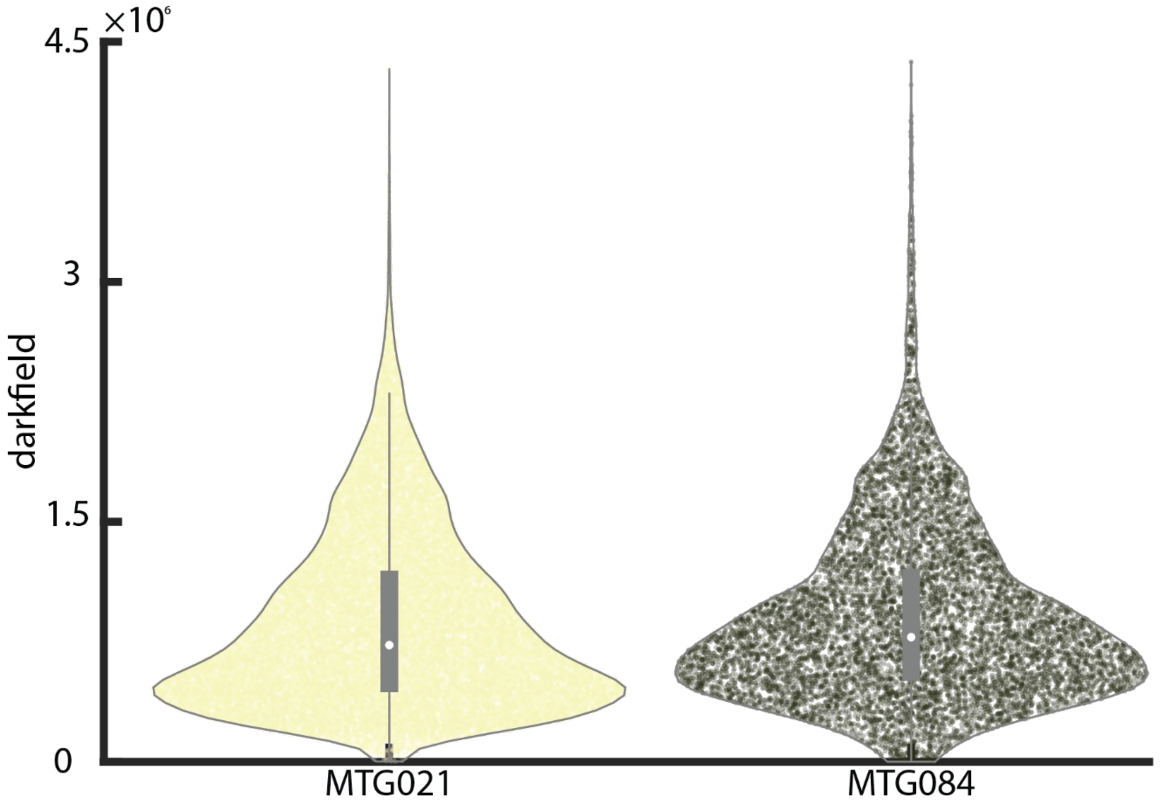


Fig. S5 Distribution of darkfield signal from MTG021 and MTG084 cell lines.

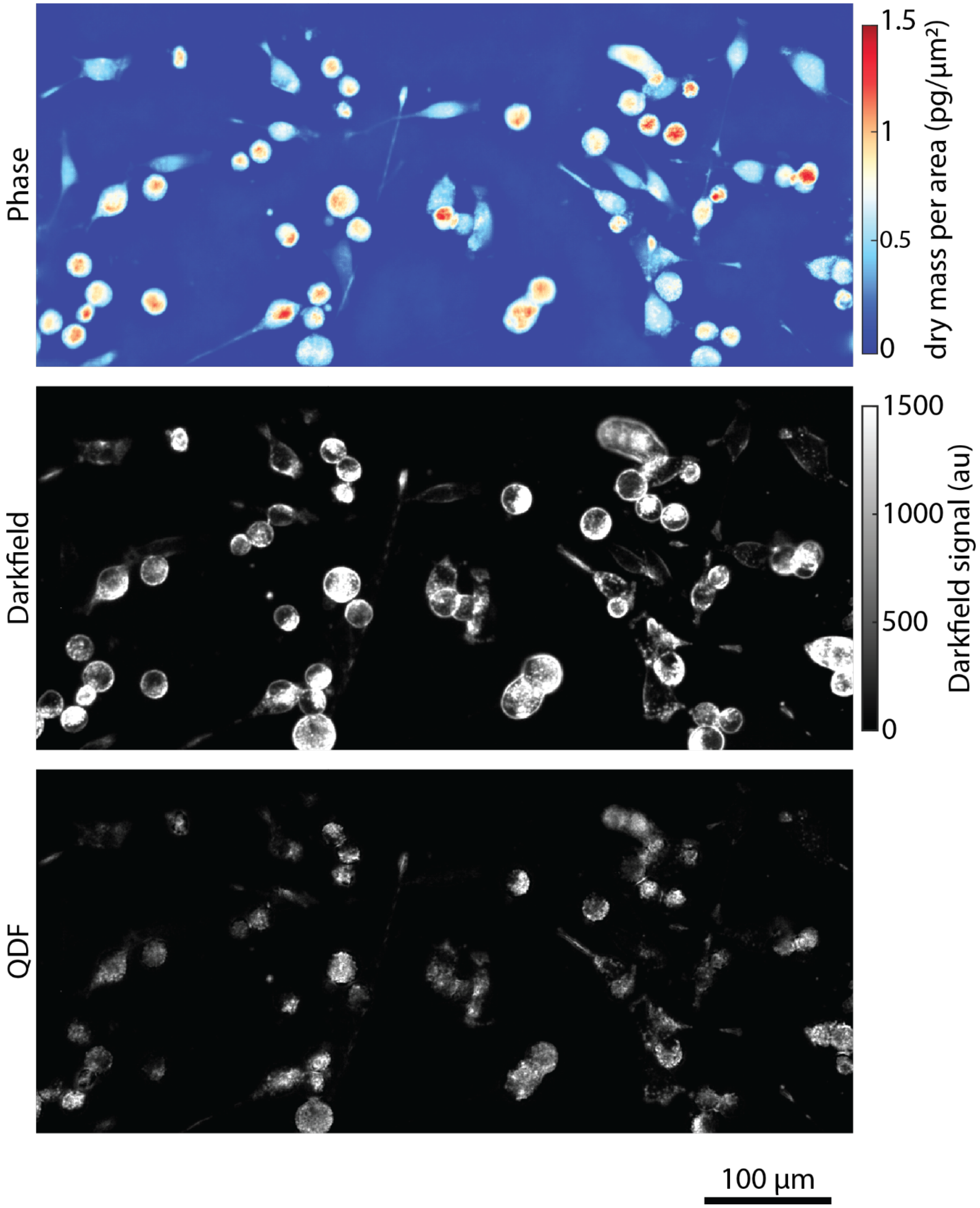


Fig. S6 Phase, Darkfield and QDF images of MTG021 showing clearer localization of puncta in QDF.
